# Dendritic degeneration and altered synaptic innervation of a central auditory neuron during age-related hearing loss

**DOI:** 10.1101/2022.11.14.516517

**Authors:** Meijian Wang, Chuangeng Zhang, Shengyin Lin, Ruili Xie

**Affiliations:** Department of Otolaryngology, The Ohio State University, Columbus, OH; Department of Neuroscience, The Ohio State University, Columbus, OH

**Keywords:** dendritic degeneration, bushy neurons, auditory nerve synapse, cochlear nucleus, age-related hearing loss, calretinin

## Abstract

Cellular morphology and synaptic configuration are key determinants of neuronal function and are often modified under pathological conditions. In the first nucleus of the central auditory system, the cochlear nucleus (CN), principal bushy neurons specialize in processing temporal information of sound critical for hearing. These neurons alter their physiological properties during aging that contribute to age-related hearing loss (ARHL). The structural basis of such changes remains unclear, especially age-related modifications in their dendritic morphology and the innervating auditory nerve (AN) synapses. Using young (2-5 months) and aged (28-33 months) CBA/CaJ mice of either sex, we filled individual bushy neurons with fluorescent dye in acute brain slices to characterize their dendritic morphology, followed by immunostaining against vesicular glutamate transporter 1 (VGluT1) and calretinin (CR) to identify innervating AN synapses. We found that dendritic morphology of aged bushy neurons had significantly reduced complexity, suggesting age-dependent dendritic degeneration, especially in neurons with predominantly non-CR-expressing synapses on the soma. These dendrites were innervated by AN bouton synapses, which were predominantly non-CR-expressing in young mice but had increased proportion of CR-expressing synapses in old mice. While somatic AN synapses degenerated substantially with age, as quantified by VGluT1-labeled puncta volume, no significant difference was observed in the total volume of dendritic synapses between young and old mice. Consequently, synaptic density on dendrites was significantly higher in old mice. The findings suggest that dendritic degeneration and altered synaptic innervation in bushy neurons during aging may underlie their changed physiological activity and contribute to the development of ARHL.

**HIGHLIGHTS:**

1. Dendrites of bushy neurons degenerate during aging with reduced complexity
2. Bushy dendrites receive AN input primarily from SGNs with medium/high threshold
3. Dendritic synapses show similar total volume but increased density during aging
4. Dendritic degeneration in bushy neurons and altered AN input may underlie ARHL

## INTRODUCTION

Sound information is transmitted from the cochlea to neurons of the cochlear nucleus (CN) via auditory nerve (AN) synapses (Nayagam et al., 2011). As the first neural station of the central auditory system, CN processes the information and projects directly or indirectly to all higher auditory nuclei to produce sound perception (Cant and Benson, 2003). Of particular interest, CN principal bushy neurons are specialized in encoding the temporal fine structure of sound (Joris et al., 1994b) critical for auditory tasks like sound localization and speech recognition (Shannon et al., 1995; Lorenzi et al., 2006; Joris and Yin, 2007; Moore, 2008; Shofner, 2008). These neurons are innervated by large somatic AN synapses called the endbulb of Held (Ryugo and Fekete, 1982; Limb and Ryugo, 2000), ensuring reliable signal transmission from the AN with temporal precision (Manis et al., 2011). We recently found that individual bushy neurons receive different subtypes of endbulbs at various convergence ratios, which correlate with their intrinsic and AN-evoked firing properties (Wang et al., 2021). AN synapses that express calretinin (CR) are from spiral ganglion neurons (SGNs) with high spontaneous rate and low threshold (type I_a_), while those without CR-expression are from SGNs with medium/low spontaneous rate and medium/high threshold (non-type I_a_) (Liberman, 1982; Petitpre et al., 2018; Sharma et al., 2018; Shrestha et al., 2018; Sun et al., 2018). The convergent innervation of type I_a_ and non-type I_a_ endbulbs on individual bushy neurons suggest that these cells play significant roles in integrating information from different sounds (e.g. different intensities) before projecting to higher nuclei. Notably, bushy neurons are also renowned for their bush-like tufted dendrites (Brawer et al., 1974; Cant and Morest, 1979; Tolbert et al., 1982; Rouiller and Ryugo, 1984), which receive only small AN bouton synapses (Gomez-Nieto and Rubio, 2009, 2011). The comprehensive innervation profile of AN synapses on the dendrites and the subtype-specificity of these synapses remain unknown. Identification of these features is crucial to understand the potential function of bushy cell dendrites.

During aging, the auditory system undergoes significant structural and functional changes leading to reduced sensory input from the cochlea and altered central auditory processing, which contribute to age-related hearing loss (ARHL) (Caspary and Llano, 2019; Xie et al., 2019; Wu et al., 2020). Particularly, synaptic transmission at AN synapses deteriorate with age (Xie and Manis, 2017a), accompanied by decreased firing rate and compromised temporal precision in postsynaptic bushy neurons (Xie, 2016). Both bushy cell body and the endbulb of Held synapses reduce in size that parallel the progress of hearing loss (McGuire et al., 2015; Connelly et al., 2017). In CBA/CaJ mice, synaptic degeneration is more severe in non-type I_a_ endbulbs during ARHL, along with a reduced prevalence of bushy neurons predominantly innervated by these synapses (Wang et al., 2021). Little is known about the age-related changes in the dendritic morphology of bushy neurons and the innervation profile of different AN synapses on the dendrites.

In this study, we investigated the morphological features of the CN bushy neurons and their innervating AN synapses in young and aged CBA/CaJ mice. We found that bushy cell dendrites degenerate during aging, especially in neurons with predominantly non-type I_a_ synapses on the soma. AN synapses on bushy cell dendrites are primarily non-type I_a_ synapse in young mice, signifying a dedicated role in processing high intensity sounds. Such preference toward non-type I_a_ synapses is lost in aged mice. These findings provide new insights in understanding the function of bushy cell dendrites and the innervating AN synapses, and how changes of both during aging may contribute to the development of ARHL.

## MATERIALS AND METHODS

CBA/CaJ mice were acquired from the Jackson Laboratory, bred and maintained at the animal facility at The Ohio State University. Two age groups of mice were used, including a young mice group with normal hearing at the age of 2 to 5 months, and an old mice group with ARHL at the age of 28 to 32 months. Experiments were performed under the guidelines of the protocol approved by the Institutional Animal Care and Use Committees at The Ohio State University (IACUC protocol # 2018A00000055-R1). Data presented in this study includes some of the bushy neurons from our previously published research (Wang et al., 2021; Zhang et al., 2022).

### Brain slice preparation

Two acute brain slices containing the CN were prepared from each mouse. As previously reported (Xie, 2016; Xie and Manis, 2017a), mice were anesthetized by intraperitoneal injection of ketamine (100 mg/kg) and xylazine (10 mg/kg) until unresponsive to toe pinch. Mice were then decapitated and skull opened to extract the brainstem, which were cut into halves at the midline and trimmed to appropriate block shape. Parasagittal slices containing the CN were cut at a thickness of 225 μm using a Vibratome 1000 (Technical Products, Inc.) or a VT1200S Microtome (Leica Biosystems, Wetzlar, Germany). The dissection and slicing procedure were performed in artificial cerebral spinal fluid (ACSF) contained (in mM): 122 NaCl, 3 KCl, 1.25 NaH_2_PO_4_, 25 NaHCO_3_, 20 glucose, 3 *myo*-inositol, 2 sodium pyruvate, 0.4 ascorbic acid, 2.5 CaCl_2_ and 1.5 MgSO_4_. The ACSF was gassed with 5% CO_2_ and 95% O_2_ and pre-warmed to 34 °C before use. Slices were incubated in the same ACSF for ∼ 45 minutes to recover before experiments began.

### Whole-cell patch clamp recording and dye-filling

Recovered slices were transferred to a recording chamber under an Axio Examiner microscope (Carl Zeiss Microscopy, LLC) and bathed in continuous flow of ACSF. The chamber was heated to 34 °C and maintained throughout the recording session. Whole-cell patch clamp recording under either current clamp or voltage clamp mode was performed from neurons in the middle- and high-frequency region of the anteroventral CN. Recording was made using Multiclamp 700B amplifier, Axon Digidata 1550B digitizer, and pClamp 10 software (Molecular Devices). Electrode pipette was fabricated from KG-33 borosilicate glass (King Precision Glass) using a Sutter P-2000 puller (Sutter Instruments). For current clamp recording, the pipette solution contained (in mM): 105 K-gluconate, 36 KCl, 2 NaCl, 10 HEPES, 0.2 EGTA, 4 MgATP, 0.3 GTP, and 10 Tris-phosphocreatine, with pH adjusted to 7.2. For voltage clamp recording, the pipette solution contained (in mM): 130 CsMetSO3, 5 CsCl, 5 EGTA, 10 HEPES, 4 MgATP, 0.3 GTP, 10 Tris-phosphocreatine, 3 QX 314, and pH adjusted to 7.20. Alexa Fluor 488 or 594 was added to the pipette solution (0.01% by weight) to fill the recorded neurons. Upon completion of data acquisition, recording pipette was slowly withdrawn from the target neuron, which resealed in most cases with the fluorescent dye trapped in the cytoplasm to allow identification and acquisition of cellular morphology. Only bushy neurons were included, which were identified based on morphological and electrophysiological properties as described previously (Brawer et al., 1974; Cant and Morest, 1979; Wu and Oertel, 1984; Xie and Manis, 2013, 2017b). Electrophysiological properties of the AN synapses and postsynaptic bushy neurons were reported in our previous publications (Wang et al., 2021; Zhang et al., 2022) and are not presented in this study.

### Post hoc immunohistochemistry

After whole-cell recording, brain slices with filled neurons were immediately fixed in 4% paraformaldehyde in PBS for 15 minutes, followed by PBS wash for 3 times at 15 minutes each. Slices were then immune-stained as previously described (Lin and Xie, 2019; Wang et al., 2021), using primary antibodies against vesicular glutamate transporter 1 (VGluT1) (polyclonal Guinea pig anti-vGluT1; Cat#: 135304, Synaptic Systems, 1:500) and calretinin (rabbit ant-CR; Cat# 214102, Synaptic Systems, 1:500). Corresponding secondary antibodies were used including goat anti rabbit IgG, Alexa 647 conjugated (Cat# A21245, Thermo Fisher Scientific, 1:500), goat anti rabbit IgG, Alexa 750 conjugated (Cat# A21039, Thermo Fisher Scientific, 1:500), goat anti guinea pig IgG, Alexa 488 conjugated (Cat # A11073, Thermo Fisher Scientific, 1:500), or goat anti guinea pig IgG, Alexa 647 conjugated (Cat # A11450, Thermo Fisher Scientific, 1:500). Stained slices were mounted on slide using DAPI-Fluoromount-G mounting media (Southern Biotech), which stained cell nuclei and helped identify the location of neighboring cells. Morphology of the labeled bushy neuron and AN synapses were acquired using an Olympus FV3000 confocal microscope (Olympus Corporation) at channels with excitation wavelength of 360, 488, 594, 647 or 750 nm. High resolution image stacks were acquired using a 60x oil-immersion objective and 2.0 – 3.0x digital zoom at a z-step size of 0.3 μm, ensuring accurate identification and measurement of synaptic and cellular structure.

### Image analysis

Image processing and measurement were performed using Imaris software (version 9.5.0; Oxford Instruments). Bushy cell morphology including the soma and dendrites were reconstructed in 3D from high resolution confocal image stacks using semi-automated Imaris tools (Wang et al., 2021). To quantify the morphological complexity of the dendrites, we performed the Sholl analysis in each bushy neuron by counting the number of dendritic intersections at fixed distances from the center point of the soma in concentric spheres. The total length of dendrites was also calculated from each bushy neuron as the summation distance that all dendritic branches traveled in space. Dendrites were further divided into primary dendrites and distal dendrites, which correspond to the portion between the soma and the first branching point as well as the portion beyond the first branching point (Fig. 1), respectively.

**Figure 1.**
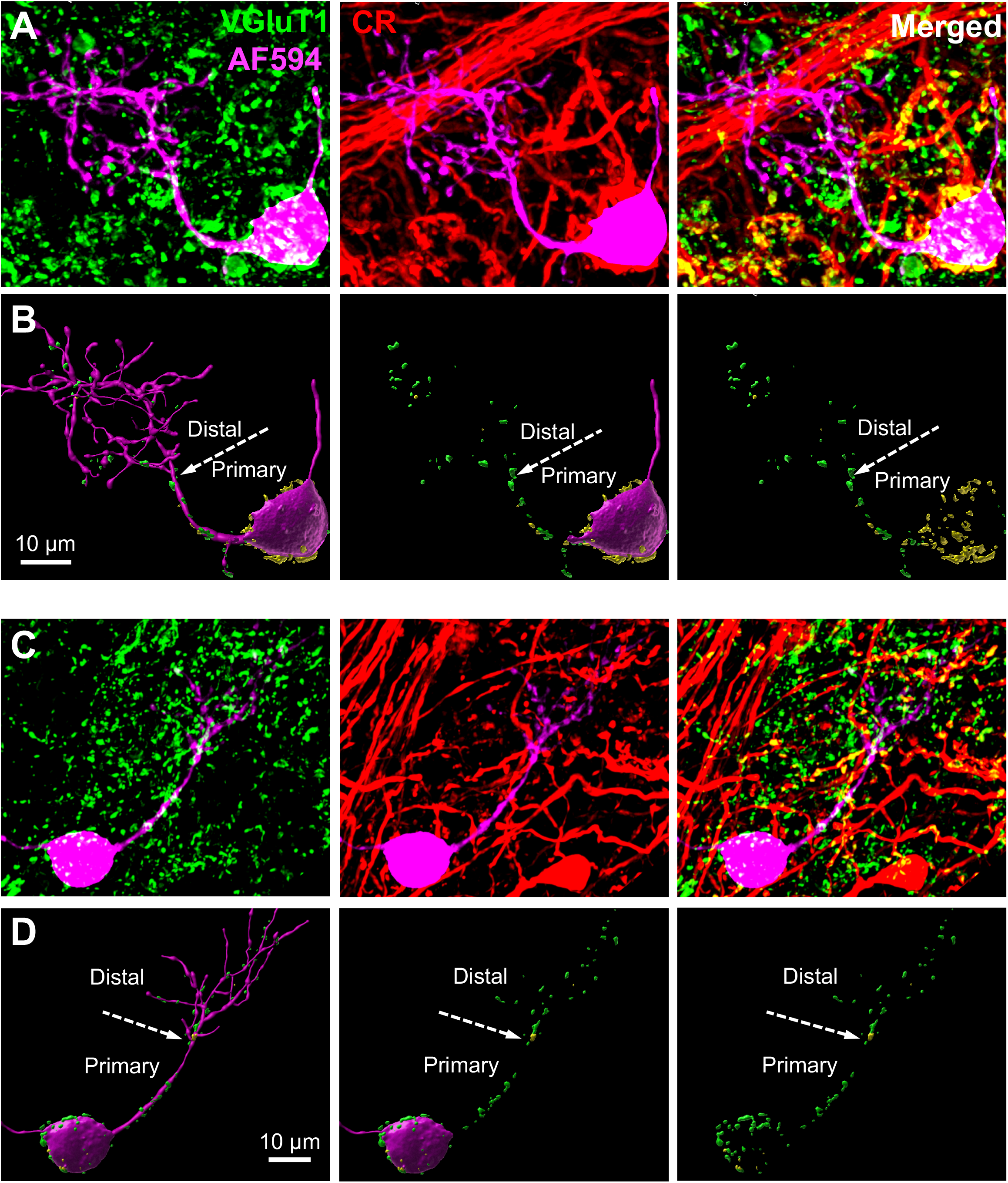
Morphology of example bushy neurons and innervating AN synapses.(**A**) Example confocal images of an example bushy neuron filled with Alexa Fluor 594 (magenta) and stained with synaptic marker VGluT1 (green) and calretinin (CR; red). In the merged panel, VGluT1-labeled puncta inside the CR^+^ terminals are shown in yellow, representing CR-expressing synapses (type I_a_); and those outside the CR^+^ terminals are shown in green, representing non-CR-expressing synapses (non-type I_a_). (B) Reconstructed bushy neuron and the innervating AN synapses from **A**. Panels show the complete cellular morphology and synapses (left), with removed dendrites (middle), and with only the synapses (right). Yellow puncta: CR-expressing synapses; green puncta: non-CR-expressing synapses. Note that this neuron receives predominantly CR-expressing synapses on the soma and predominantly non-CR-expressing synapses on the dendrites. Arrows mark the first branching point that divides the primary and distal dendrites. (**C-D**) Another example bushy neuron that receives predominantly non-CR-expressing synapses on both soma and dendrites.

AN synaptic input on the cell body (somatic synapses) and dendrites (dendritic synapses) of the target bushy neuron were also reconstructed and their subtype-specificity identified based on the patterns of VGluT1- and CR-staining (Wang et al., 2021). Specifically, type I_a_ AN synapses were labeled by both VGluT1- and CR-staining, while non-type I_a_ AN synapses were only labeled by VGluT1-, but not by CR-staining. Consistent with previous studies (Wang et al., 2021), we quantified the volume of VGluT1-labeled puncta from AN synapses on the soma of every bushy neuron and classified all neurons into different groups. The puncta volume of AN bouton synapses on the dendrites of each bushy neuron were also measured.

### Statistical analysis

Statistical analyses were performed using GraphPad Prism (Version 6.0h, GraphPad Software). Group data were first tested for normality (D’Agostino & Pearson omnibus normality test) to determine if they were normally distributed, followed by unpaired t-test (normal distribution) or Mann-Whitney test (non-normal distribution). Two-way ANOVA was also used to examine the effect of different cell types and ages on bushy cell morphology and AN synaptic input. Data are presented as mean ± standard deviation.

## RESULTS

Data in this study were acquired from 37 young mice at the age of 2-5 months, and 33 old mice at the age of 28-32 months. Hearing status of these mice were assessed by measuring their auditory brainstem response, which respectively showed normal hearing and severe ARHL as previously reported (Wang et al., 2021).

### Bushy cell morphology and the innervating AN synapses

In order to investigate the cellular morphology of bushy neurons and the innervating AN synapses, we performed whole-cell recording from these cells in CN slices, filled individual neurons with Alexa Fluor 594, followed by post hoc immunostaining using antibodies against VGluT1 and CR (see Materials and Methods for details). Electrophysiological properties of the bushy neurons and AN synapses were reported previously (Wang et al., 2021; Zhang et al., 2022) and the present study only focused on morphology. As shown in Fig. 1, bushy neurons had typically one primary dendrite and numerous fine branches at the distal end. Staining of the synaptic marker VGluT1 showed that bushy neurons not only receive large synaptic inputs on the soma, but also small bouton synapses on both primary and distal dendrites (Fig. 1B, D). Based on the localization of VGluT1 and CR labeling, we were able to identify the CR-expressing (CR, or type I_a_) and non-CR-expressing (non-CR, or non-type I_a_) synapses, which were shown in yellow and green in the merged panel (Fig. 1A, C) (Wang et al., 2021). We acquired image stacks of the target cells using confocal microscope and reconstructed the 3D structure of the bushy neurons and their innervating AN synapses (Fig. 1B, D). Bushy cell morphology, AN synapses on bushy cell soma (Wang et al., 2021; Zhang et al., 2022) and dendrites were analyzed in neurons from young and old mice to evaluate morphological changes during ARHL.

### Dendritic morphology is less complex in bushy neurons from old mice

We first assessed the dendritic morphology of bushy neurons from normal hearing young mice and old mice with ARHL. As shown in Fig. 2A, bushy neurons from young mice had extensive dendritic trees with numerous distal branches. In old mice, some bushy neurons retained similar dendritic morphology as exampled by the top two neurons of the old group. Other neurons, however, showed simplified dendritic morphology with shorter and fewer number of distal branches. To quantify the complexity of the dendritic morphology among these neurons, we performed the Sholl analysis in each cell by counting the number of dendritic intersections at fixed distances from the soma. On average, bushy neurons in young mice had significantly more dendrite intersections than those in old mice (Fig. 2B; Two-way ANOVA: age effect, *F*_*1, 1539*_ = 48.7, *P* < 0.0001; distance effect, *F*_*18, 1539*_ = 38.3, *P* < 0.0001; interaction, *F*_*18, 1539*_ = 4.04, *P* < 0.0001). We further traced all dendritic branches and calculated the total dendrite length, which was significantly longer in bushy neurons from the young mice than those from the old mice (Fig. 2C; total dendritic length in young: 477 ± 193 μm, n = 47; old: 299 ± 227 μm, n = 41; Mann Whitney test: *P* < 0.0001). Specifically, the length of primary dendrites was not changed (young: 35 ± 13 μm; old: 35 ± 16 μm; Mann Whitney test: *P* = 0.756), whereas the total length of distal dendrites was significantly reduced (young: 447 ± 196 μm; old: 292 ± 225 μm; Mann Whitney test: *P* = 0.0007). We also measured the volume of the cell body, which was significantly smaller in old mice (Fig. 2D; young: 1005 ± 331 μm^3^; old: 681 ± 245 μm^3^; Mann Whitney test: *P* < 0.0001). The results indicate that during aging, bushy neurons degenerate in distal dendrites and reduce in cell body size.

**Figure 2.**
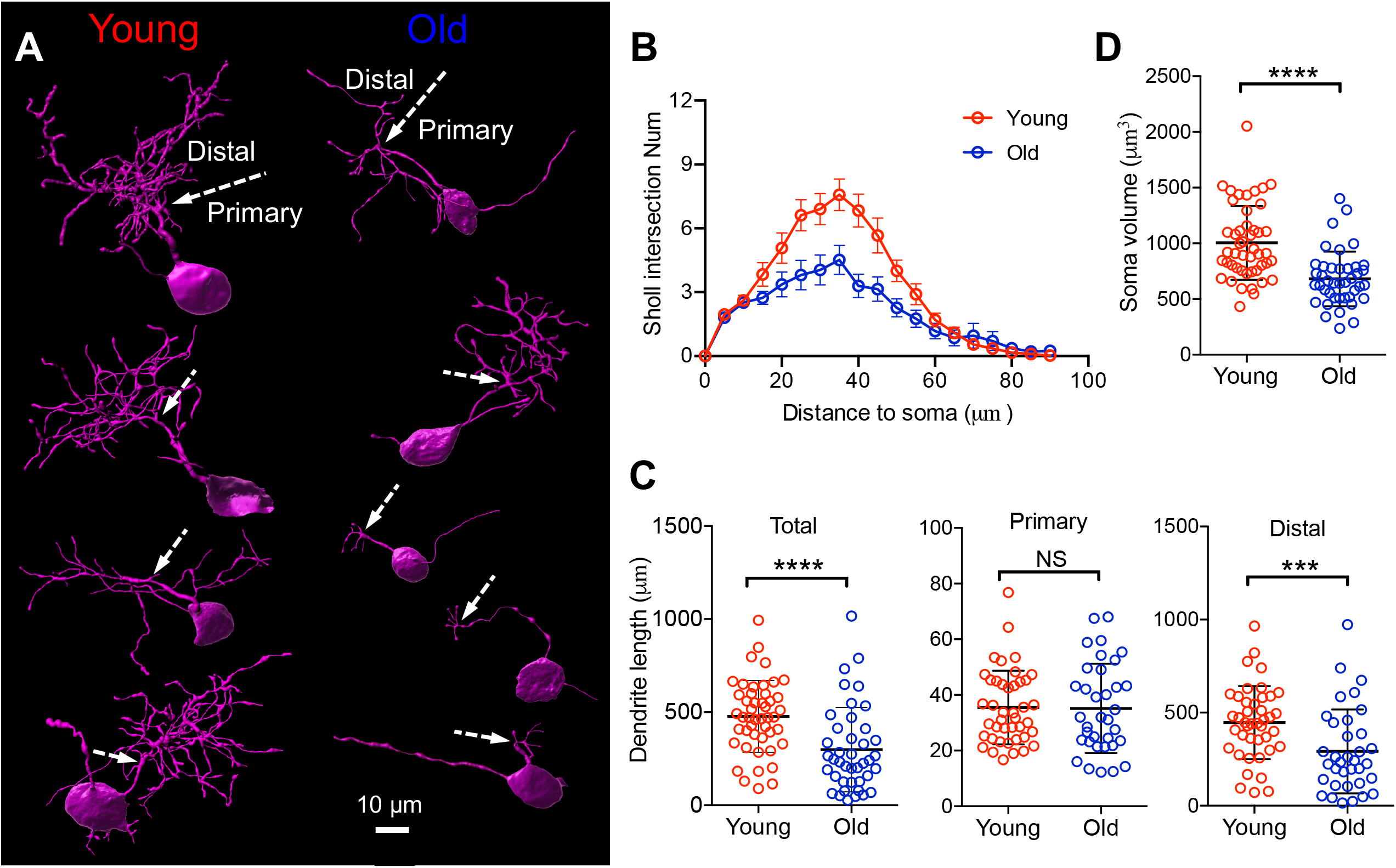
Bushy neurons show degenerated distal dendrites with age.(**A**) Example reconstructed bushy neurons from young and old mice. Arrows mark the first branching points that divide the primary from distal dendrites. (**B**) Sholl analysis of young and old bushy neurons to show reduced dendritic morphology. (**C**) Summary of total dendritic length, primary dendritic length, and distal dendritic length of all bushy neurons. (**D**) Summary of soma volume. NS, not significant; \*\*\**P* < 0.001; \*\*\*\**P* < 0.0001.

### Dendritic degeneration is more profound in neurons with predominantly non-CR-expressing synapses on the soma

Our previous studies showed that bushy neurons with different subtypes of AN synapses on the soma exhibit distinct intrinsic properties and AN-evoked responses (Wang et al., 2021; Zhang et al., 2022). It is unclear whether these bushy neurons differ in their dendritic morphology and how the morphology might differentially change during aging. To answer these questions, we classified bushy neurons into two subgroups using the same approach as in our previous studies (Wang et al., 2021; Zhang et al., 2022), by identifying the subtype-specificity of AN synapses on the soma and measuring their VGluT1-labeled puncta volume. Bushy neurons with predominantly CR-expressing synapses on the soma (> 60% CR-expressing synapses by volume) were classified as CR-dominant neurons (Fig. 3A, left column), and those with predominantly non-CR-expressing synapses on the soma (< 40% CR-expressing synapses by volume) were classified as non-CR-dominant neurons (Fig. 3A, right column). In young mice, Sholl analysis of the dendritic trees showed no significant difference between two subgroups of bushy neurons (Fig. 3B; Two-way ANOVA: cell type effect, *F*_*1, 798*_ = 1.98, *P* = 0.1602; distance effect, *F*_*18, 798*_ = 27.8, *P* < 0.0001; interaction, *F*_*18, 798*_ = 1.12, *P* = 0.3252). Their total dendrite length was also similar (Fig. 3C; CR-dominant neurons: 501 ± 148 μm, n = 24; non-CR-dominant neurons: 458 ± 238 μm, n = 20; Mann-Whitney test: *P* = 0.3555). It suggests that despite having different physiological properties (Wang et al., 2021; Zhang et al., 2022), bushy neurons with predominantly CR- and non-CR-expressing AN synapses on the soma have similar dendritic morphology.

**Figure 3.**
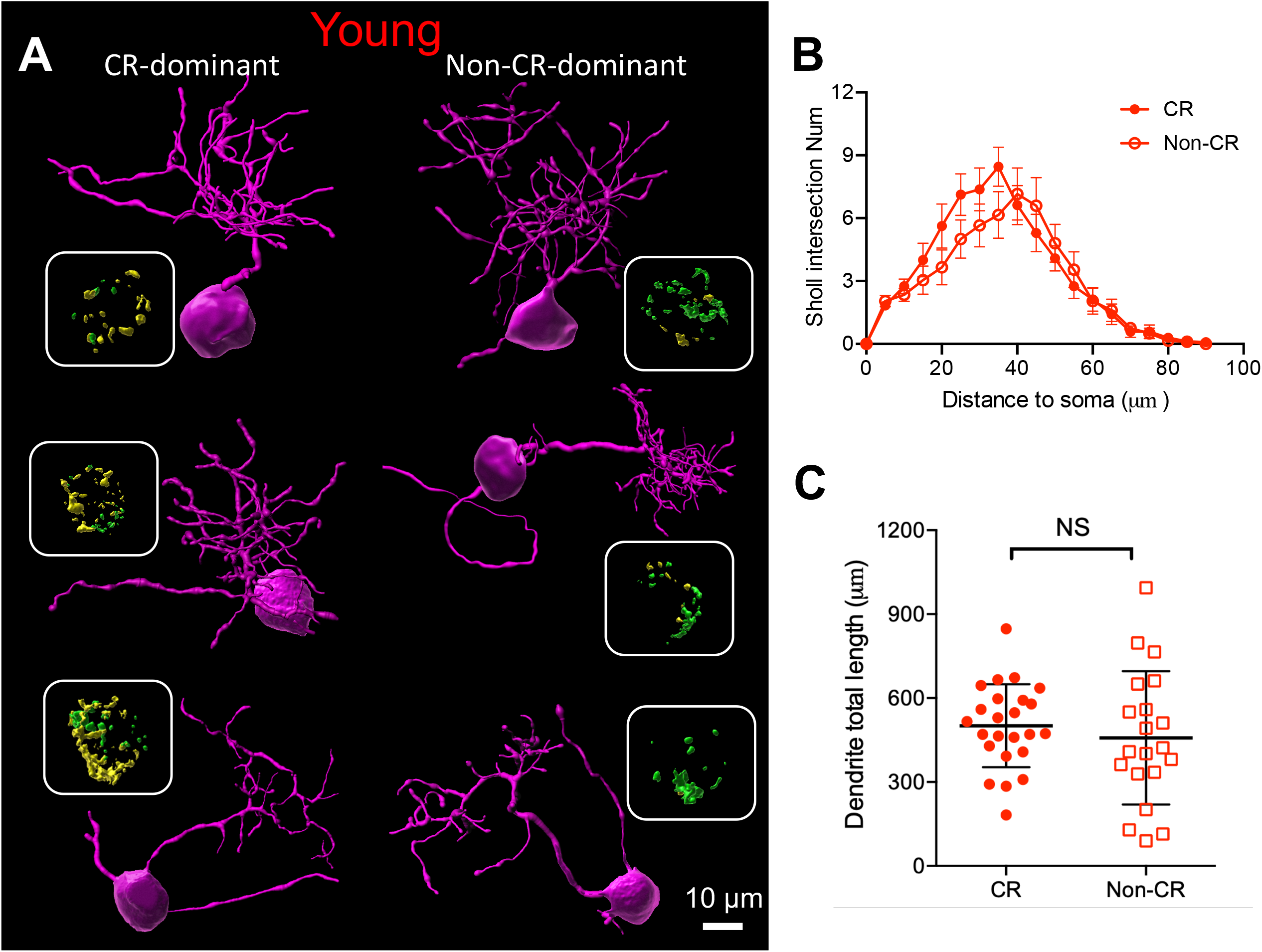
Bushy neurons with different subtypes of somatic synapses have similar dendritic morphology in young mice.(**A**) Reconstructed bushy neurons with predominantly CR-expressing (left column) and non-CR-expressing (right column) synapses on the soma. Insets show reconstructed somatic synapses. Yellow: CR-expressing synapses; green: non-CR-expressing synapses. (**B**) Sholl analysis of the dendritic morphology in bushy neurons with CR- and non-CR-dominant synapses. (**C**) Summary of total dendritic length. NS, not significant.

We performed the same analysis in bushy neurons from old mice with ARHL. The dendritic morphology of non-CR-dominant bushy neurons was less complex than those of CR-dominant neurons (Fig. 4A), with significantly less dendrite intersections (Fig. 4B; Two-way ANOVA: cell type effect, *F*_*1, 741*_ = 24.1, *P* < 0.0001; distance effect, *F*_*18, 741*_ = 10.2, *P* < 0.0001; interaction, *F*_*18, 741*_ = 2.31, *P* = 0.0016) and shorter total dendrite length (Fig. 4C; CR neurons: 381 ± 242 μm, n = 22; non-CR neurons: 203 ± 167 μm, n = 19; Mann-Whitney test: *P* = 0.0070). Therefore, while dendrites degenerate with age in both subgroups of bushy neurons, the degeneration is more severe in non-CR-dominant cells.

**Figure 4.**
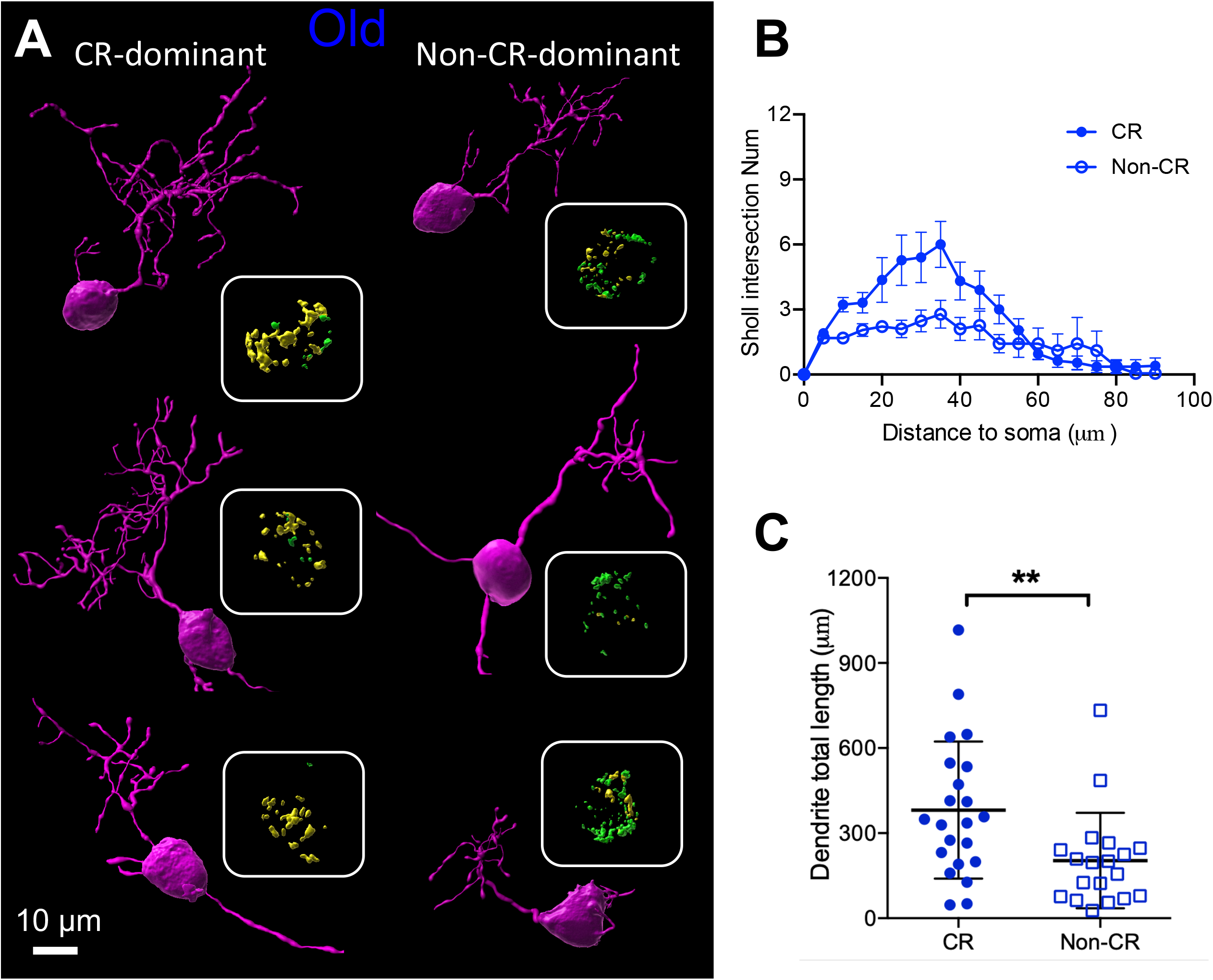
Bushy neurons with non-CR-dominant synapses on the soma show less complex dendritic morphology in old mice.(**A**) Reconstructed bushy neurons with predominantly CR-expressing (left column) and non-CR-expressing (right column) synapses on the soma. Insets show reconstructed somatic synapses. Yellow: CR-expressing synapses; green: non-CR-expressing synapses. (**B**) Sholl analysis of the dendritic morphology in bushy neurons with CR- and non-CR-dominant synapses. (**C**) Summary of total dendritic length. \*\**P* < 0.01.

### Innervation profile of AN synapses on bushy neurons and changes during aging

Bushy neurons are known to receive large endbulb of Held synapses on the soma (Ryugo and Fekete, 1982; Manis et al., 2011), and small bouton synapses on the dendrites (Gomez-Nieto and Rubio, 2009, 2011). However, the comprehensive innervation profile of AN synapses on bushy neurons remains unclear, especially regarding the subtype specificity of the synapses and their changes during aging. As shown in Fig. 1, we reconstructed the somatic and dendritic synapses on bushy neurons and quantified these synapses by measuring their VGluT1-labeled puncta volume. In young mice, the total volume of somatic synapses was significantly larger on average than that of the dendritic synapses (Fig. 5A; somatic synapses: 61 ± 36 μm^3^; dendritic synapses: 16 ± 12 μm^3^; Wilcoxon matched-pairs signed rank test: *P* < 0.0001). It suggests that AN inputs onto bushy neurons are dominated by somatic synapses, especially after considering the attenuation of dendritic signals due to filtering effect of dendrites (Spruston et al., 1994). We further calculated the volume percentage of CR-expressing synapses on the soma and dendrites and plotted in Fig. 5B. The volume percentage of CR-expressing synapses on the soma varied on a continuum along the x-axis (left panel), consistent with our previous findings (Wang et al., 2021). However, a majority of bushy neurons (34/43, right panel) received predominantly non-CR-expressing synapses on the dendrites (CR% < 50%). As non-CR-expressing synapses correspond to SGNs with medium/low spontaneous rate and medium/high threshold (Liberman, 1982; Petitpre et al., 2018; Sharma et al., 2018; Shrestha et al., 2018; Sun et al., 2018), it suggests that the dendrites of bushy neurons may serve as an dedicated region to preferentially process auditory information from medium/loud intensity sounds. Interestingly, there was a significant correlation that bushy neurons with higher percentage of CR-expressing synapses on the soma receive higher percentage of CR-expressing synapses on the dendrites (Fig. 5B left panel; linear regression: *r*^2^ = 0.214, *P* = 0.0018).

**Figure 5.**
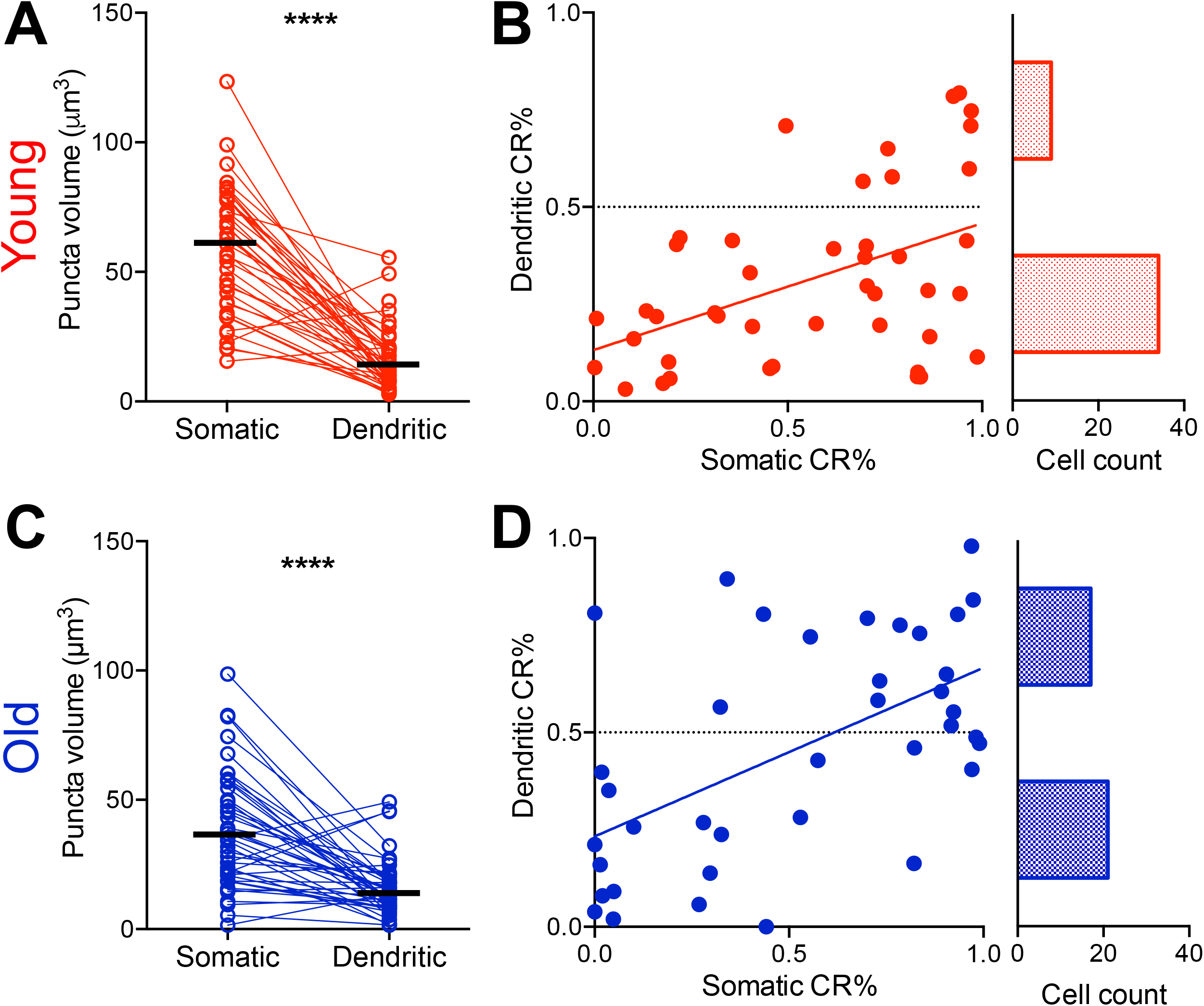
Changes of AN synapses on bushy neurons during aging.(**A**) Volume of somatic and dendritic synapses onto bushy neurons in young mice. Connected circles are from the same neurons. (**B**) Left panel: percentage of CR-expressing synapses on the soma (x-axis) and dendrites (y-axis). Red line: linear regression, *r*^2^ = 0.214, *P* = 0.0018. Right panel: cell count of bushy neurons with predominant CR-expressing (n = 9; CR% > 50%) and non-CR-expressing (n = 34; CR% < 50%) synapses on dendrites. (**C**)Volume of somatic and dendritic synapses onto bushy neurons in old mice. (**D**) Left panel: percentage of CR-expressing synapses on the soma and dendrites. Blue line: linear regression, *r*^2^ = 0.313, *P* = 0.0003. Right panel: cell count of bushy neurons with predominant CR-expressing (n = 17; CR% > 50%) and non-CR-expressing (n = 21; CR% < 50%) synapses on dendrites. \*\*\*\**P* < 0.0001.

In old bushy neurons, the volume of somatic synapses remains significantly larger than that of dendritic synapses (Fig. 5C; somatic synapses: 38 ± 22 μm^3^; dendritic synapses: 17 ± 12 μm^3^; Wilcoxon matched-pairs signed rank test: *P* < 0.0001). Compare to young bushy neurons, old bushy neurons had smaller somatic synapses by volume (Fig. 5 A, C; Mann Whitney test: *P* = 0.0002), consistent with our previous report of synaptic degeneration during aging (Wang et al., 2021). Surprisingly, there was no significant difference in the volume of dendritic synapses between young and old bushy neurons (Fig. 5A, C; Mann Whitney test: *P* = 0.6278). Furthermore, there were about equal numbers of bushy neurons that received predominantly CR-expressing (17/38; CR% > 50%) and non-CR-expressing (21/38; CR% < 50%) synapses on dendrites (Fig. 5D, y-axis), suggesting a relative shift in subtype-specificity of dendritic synapses from non-CR-expressing to CR-expressing synapses during aging. The correlation that bushy neurons with higher percentage of CR-expressing synapses on the soma receive higher percentage of CR-expressing dendritic synapses was retained in old mice (Fig. 5D left panel; linear regression: *r*^2^ = 0.313, *P* = 0.0003).

We further investigated the changes of dendritic synapses during aging by separately analyzing the synapses on primary and distal dendrites, as exampled in Fig. 6A. On average, synaptic volume was not significantly different in total dendritic synapses, synapses on primary dendrites, or synapses on distal dendrites between young and old bushy neurons (Fig. 6B; young primary: 6.5 ± 7.2 μm^3^, old primary: 6.1 ± 7.3 μm^3^, Mann Whitney test: *P* = 0.9450; young distal: 8.7 ± 7.5 μm^3^, old distal: 10.1 ± 8.7 μm^3^, Mann Whitney test: *P* = 0.6142). Dendritic synaptic density was calculated as the VGluT1-labeled puncta volume divided by dendrite length, which significantly increased in old bushy neurons in total and distal dendrites (Fig. 6C; young total: 0.045 ± 0.054 μm^3^/μm, old total: 0.082 ± 0.069 μm^3^/μm, Mann Whitney test: *P* = 0.0020; young distal: 0.028 ± 0.039 μm^3^/μm, old distal: 0.057 ± 0.058 μm^3^/μm, Mann Whitney test: *P* = 0.0055), but not in primary dendrites (young primary: 0.20 ± 0.21 μm^3^/μm, old primary: 0.18 ± 0.14 μm^3^/μm, Mann Whitney test: *P* = 0.6520). It suggests that AN synapses on bushy cell dendrites maintain stable synaptic volume during aging but increase in synaptic density, particularly on distal dendrites that consistent with distal dendrite degeneration (Fig. 2C).

**Figure 6.**
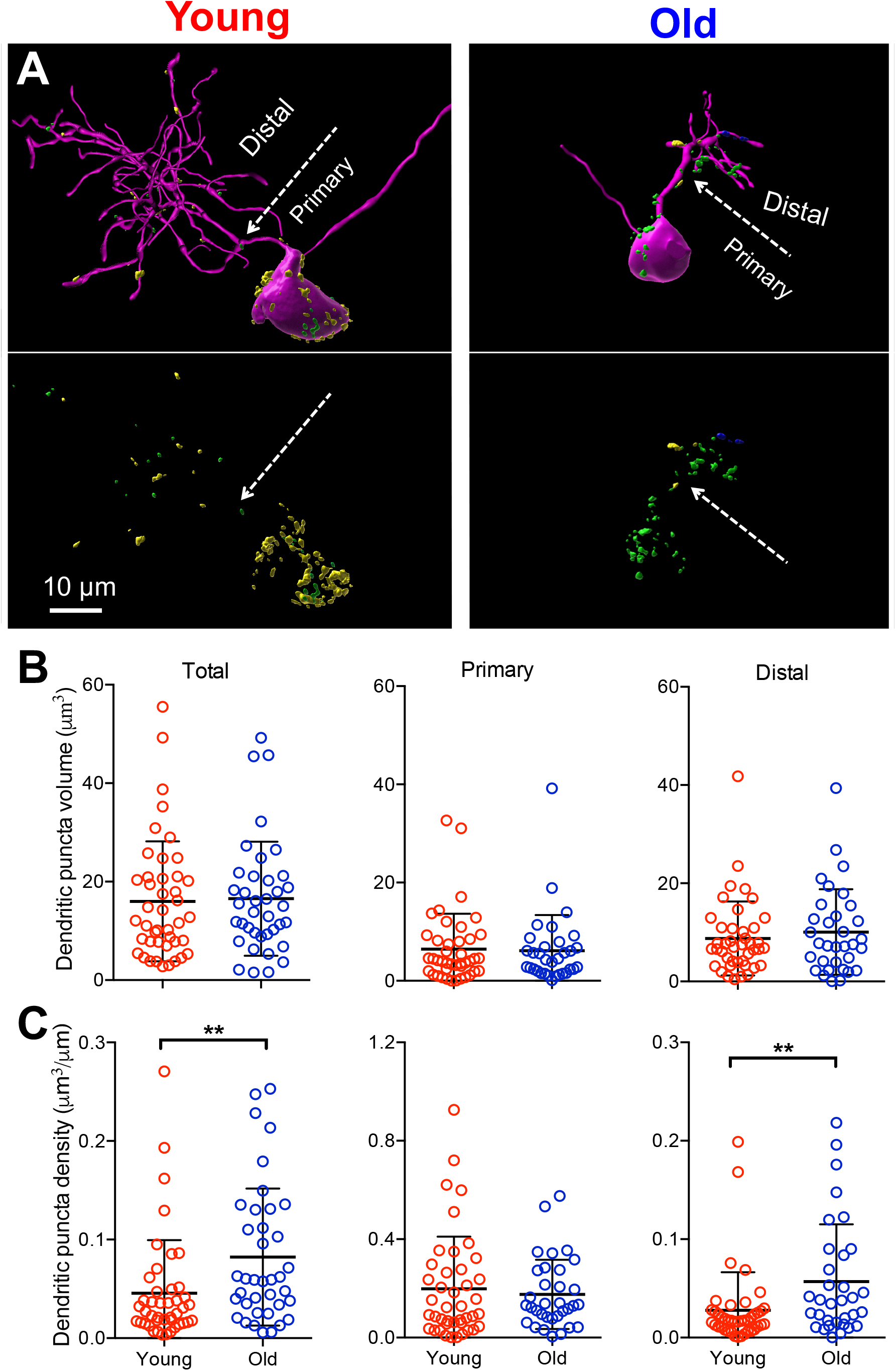
Summary changes of dendritic synapses during aging.(**A**) Example reconstructed bushy neurons and innervating AN synapses in young and old mice. Note that the old bushy neuron had less complex dendritic morphology and denser dendritic synapses. (**B**) Summary of puncta volume of total dendritic synapses, synapses on primary dendrites, and synapses on distal dendrites. (**C**) Summary of synaptic puncta density on total dendrites, primary dendrites and distal dendrites. \*\**P* < 0.01.

Finally, we studied the volume and density of the dendritic synapses from CR-dominant and non-CR-dominant bushy neurons. In young mice, no significant difference was found in either synaptic volume (Fig. 7A; CR total: 15.7 ± 9.6 μm^3^, non-CR total: 16.4 ± 15.1 μm^3^, Mann Whitney test: *P* = 0.5529; CR primary: 6.7 ± 8.6 μm^3^, non-CR primary: 6.1 ± 4.8 μm^3^, Mann Whitney test: *P* = 0.5543; CR distal: 8.9 ± 5.7 μm^3^, non-CR distal: 8.5 ± 9.6 μm^3^, Mann Whitney test: *P* = 0.2950), or synaptic density (Fig. 7B; CR total: 0.038 ± 0.039 μm^3^/μm, non-CR total: 0.055 ± 0.068 μm^3^/μm, Mann Whitney test: *P* = 0.8182; CR primary: 0.19 ± 0.24 μm^3^/μm, non-CR primary: 0.21 ± 0.18 μm^3^/μm, Mann Whitney test: *P* = 0.3574; CR distal: 0.021 ± 0.015 μm^3^/μm, non-CR distal: 0.038 ± 0.056 μm^3^/μm, Mann Whitney test: *P* = 0.7343). In old mice, there was no significant difference in dendritic synaptic volume (Fig. 7C; CR total: 19.0 ± 14.3 μm^3^, non-CR total: 13.8 ± 6.9 μm^3^, unpaired t test: *P* = 0.1702; CR primary: 7.6 ± 9.9 μm^3^, non-CR primary: 4.7 ± 3.2 μm^3^, Mann Whitney test: *P* = 0.9575; CR distal: 11.0 ± 10.9 μm^3^, non-CR distal: 9.2 ± 6.3 μm^3^, Mann Whitney test: *P* > 0.9999). However, the total synaptic density was significantly higher in non-CR-dominant neurons (Fig. 7D; CR total: 0.066 ± 0.072 μm^3^/μm, non-CR total: 0.100 ± 0.064 μm^3^/μm, Mann Whitney test: *P* = 0.0476), although no significant difference was found in either primary or distal dendrites (Fig. 7D; CR primary: 0.16 ± 0.15 μm^3^/μm, non-CR primary: 0.19 ± 0.13 μm^3^/μm, Mann Whitney test: *P* = 0.3090; CR distal: 0.037 ± 0.051 μm^3^/μm, non-CR distal: 0.075 ± 0.060 μm^3^/μm, Mann Whitney test: *P* = 0.0579). The results suggest that AN bouton synapses innervate the dendrites of different subgroups of bushy neurons similarly in terms of synaptic volume and density under normal hearing conditions. However, the synaptic density on the dendrites of non-CR-dominant neurons become higher in aged mice, which likely represents compensatory changes to dendritic degeneration (Fig. 4) and preferential damage of SGNs with medium/low spontaneous rate (Kujawa and Liberman, 2009; Furman et al., 2013; Liberman, 2017) during ARHL.

**Figure 7.**
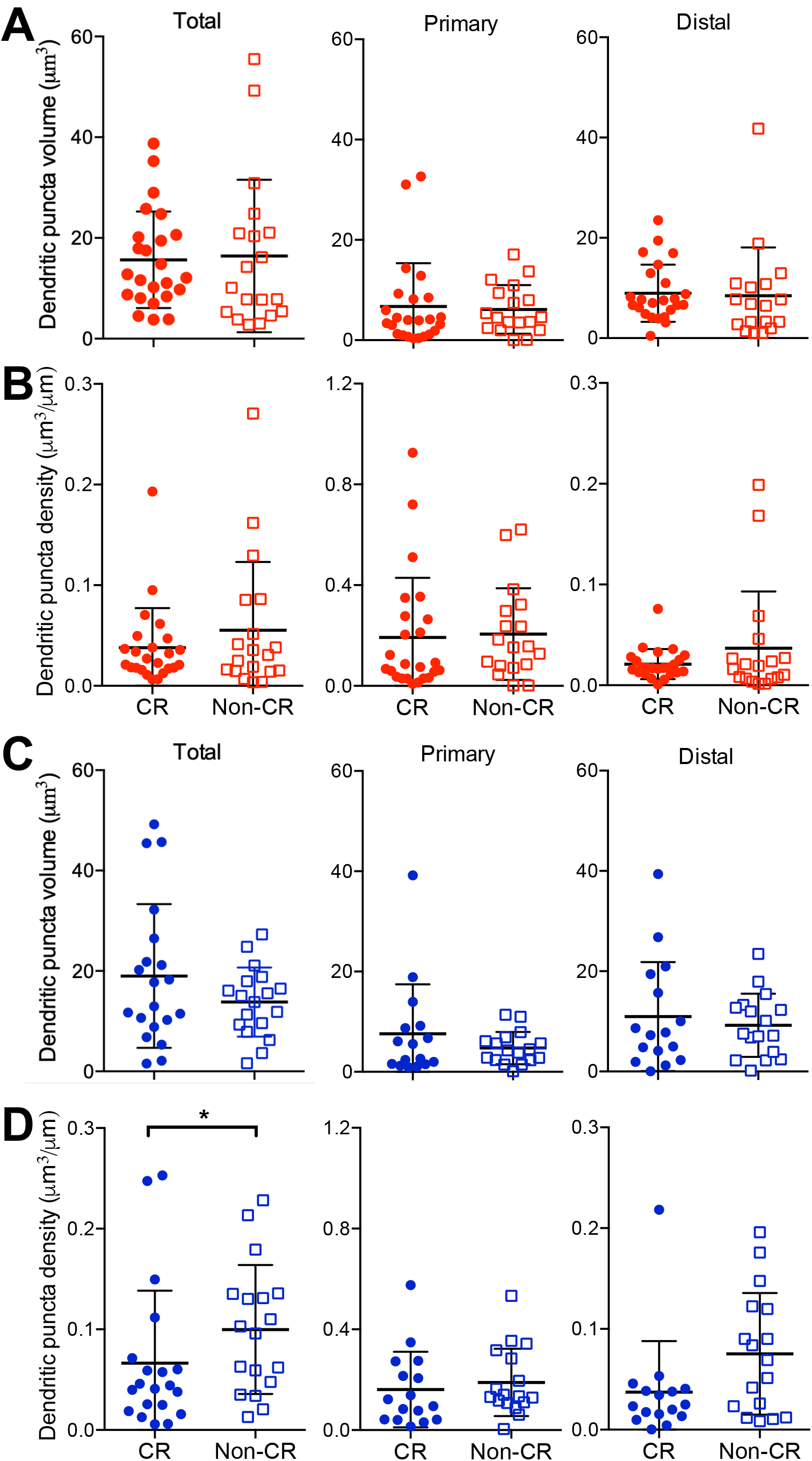
Dendritic synapses show higher puncta density in non-CR-dominant bushy neurons in old mice.(**A**) Summary of total dendritic puncta volume, volume on primary dendrites and distal dendrites between CR-and non-CR-dominant neurons in young mice. (**B**) Summary of total dendritic puncta density, puncta density on primary dendrites and distal dendrites between CR- and non-CR-dominant neurons in young mice. (**C-D**) Summary data of synaptic puncta volume and density in neurons from old mice. \**P* < 0.05.

## DISCUSSION

Bushy neurons are known for their unique bush-like cellular morphology (Brawer et al., 1974; Cant and Morest, 1979; Tolbert et al., 1982; Rouiller and Ryugo, 1984). Despite decades of research, studies that characterize both bushy cell morphology and the comprehensive innervation pattern of connecting AN synapses are rare (Spirou et al., 2022), especially regarding to AN synapses on dendrites, the subtype-specificity of AN synapses, and changes of these features during ARHL. Combining electrophysiology and immunohistochemistry, we developed the technique to investigate both the physiology and morphology of individual bushy neurons and their AN synapses in young and aged mice (Wang et al., 2021; Zhang et al., 2022). For the first time, this study characterized the changes of dendritic morphology in bushy neurons and associated AN synapses during aging, with findings that shed new light on potential mechanisms supported by such morphological features under normal hearing and ARHL.

### Function of bushy cell dendrites and degeneration during ARHL

Bushy neurons are one of the principal neurons of the CN and specialize in processing temporal information of sound (Joris et al., 1994b; Joris et al., 1994a; Manis et al., 2011), which is crucial for hearing and compromised during ARHL (Grose and Mamo, 2010; Walton, 2010; Anderson et al., 2012). Mechanisms underlying their exceptional ability of encoding auditory information with temporal precision were widely studied, including the expression of unique combination of voltage-gated ion channels (Manis and Marx, 1991; Rothman and Manis, 2003; Cao et al., 2007), fast AMPA receptors (Gardner et al., 1999; Rubio et al., 2017), low input resistance (Wu and Oertel, 1984), narrow integration window for synaptic inputs (McGinley and Oertel, 2006), as well as the somatic location of large endbulb of Held synapses (Ryugo and Fekete, 1982; Rouiller et al., 1986; Manis et al., 2011). Bushy neurons maintain an elaborate dendritic tree that is expected to contribute to their physiological function, which were under-studied but speculated as below with limited evidence.

Dendrites are generally viewed as the main site of a neuron to receive incoming synaptic inputs, which is unlikely the case in bushy neurons. Gomez-Nieto and Rubio (2009, 2011) and Spirou et al (2022) reported that bushy neurons receive small AN synapses on the dendrites, which presumably help transmit sound information from the cochlea. Our measurement showed that the total volume of dendritic synapses is only 26% of the somatic synapses on average in bushy neurons from normal hearing young mice (Fig. 5A). These dendritic synapses are probably not a major source of excitatory drive to elicit spikes in bushy neurons, especially after considering the fact that dendritic signals are substantially attenuated due to the cable property of the dendrites (White et al., 1994; Sumner et al., 2009; Koert and Kuenzel, 2021). Besides the excitatory AN synapses, inhibitory synapses were also found on the dendrites (Gomez-Nieto and Rubio, 2009, 2011; Spirou et al., 2022), which help provide modulatory inhibition to regulate discharge rate and temporal precision of bushy neurons (Caspary et al., 1994; Kopp-Scheinpflug et al., 2002; Gai and Carney, 2008; Xie and Manis, 2013; Kuenzel et al., 2015). However, the low synaptic density on dendrites (Fig. 1, 5A) (Gomez-Nieto and Rubio, 2011) and complete lack of any synapse on some branches (Spirou et al., 2022) suggest that bushy cell dendrites also contribute to other functions.

Besides receiving synapses, dendrites were also postulated to increase membrane surface to house various ion channels and pumps that help maintain intracellular ionic balance (Brownell and Manis, 2014). One particular ion is K^+^, which is constantly released from the cell through low-voltage activated K^+^ channels on the soma that needs to be replenished elsewhere. Such process increases the overall leakiness of the neuron and contributes to their unique biophysical properties. Indeed, computer model simulation showed that without altering synaptic inputs, reducing surface area by pruning dendrites is sufficient to increase membrane input resistance and time constant, and lead to enhanced neuronal excitability and reduced temporal precision (Spirou et al., 2022). Our previous study showed that bushy neurons from aged mice exhibit increased excitability and compromised temporal precision (Xie, 2016). Such physiological changes are likely contributed by the age-dependent dendritic degeneration observed in this study (Fig. 2). The findings suggest that pruning of dendrites may be an important central mechanism during aging to boost neuronal excitability and increase the responsiveness of auditory neurons, resulting in enhanced central gain to compensate for the weakened sensory input from the cochlea under ARHL.

Bushy cell dendrites may also be important to support the network activity in the ventral CN. Gomez-Nieto and Rubio reported that multiple bushy neurons form a complex neural network that are likely electrically coupled via membrane junctions on the dendrites (2009, 2011). These dendrites often orient toward the soma of neighboring cells that may help synchronize neuronal activity among different bushy neurons and promote population responses. Degeneration of bushy cell dendrites during aging is expected to disrupt such connection and reduce network activity, resulting in compromised signal processing in the CN under ARHL.

### Spatial segregation of non-type I_a_ AN synapses on bushy cell dendrites

Our study showed that bushy neurons receive different subtypes of AN synapses on the soma at a continuum of convergence ratios (Fig. 5B, x-axis), consistent with our previous report (Wang et al., 2021). AN synapses innervating the dendrites, however, are dominated by non-CR-expressing (non-type I_a_) synapses in most bushy neurons from normal hearing mice (Fig. 5B, y-axis). It suggests that bushy cell dendrites may serve as a spatially segregated compartment that facilitate the processing of auditory information from SGNs with medium/low spontaneous rate and medium/high thresholds. As discussed above, these dendritic synapses are small in size and not the main driving force to trigger spikes in bushy neurons. Nevertheless, they help modulate the efficacy of the main inputs on the soma and improve the temporal precision and fidelity of encoded information in bushy neurons (Koert and Kuenzel, 2021). Given the functional significance of SGNs with medium/low spontaneous rate in coding loud sounds (Liberman, 1978), the findings suggest that bushy cell dendrites may play unique roles in encoding specific auditory signals, especially in noisy environment. We also found that in aged mice, non-type I_a_ synapses no longer prevail in dendritic synapses among bushy neurons (Fig. 5D, y-axis), suggesting that such roles may be compromised under ARHL.

### Limitations of the study

As a cross-sectional study, we collected all the data from different mice at specific ages that do not allow assessment of longitudinal changes of individual bushy neurons and AN synapses. Our observations only reflect the status of the surviving bushy neurons during ARHL, as lost bushy neurons during aging (Willott et al., 1987) could not be sampled in our neuronal population. Such limitation in sampling may impact the interpretation of the observed results. For example, the finding that dendritic synapses maintain similar synaptic volume (Fig. 5A, C; 6B) during aging was unexpected and counterintuitive given the significant degeneration of dendrites in old bushy neurons. It is likely that the remaining AN synapses do re-organize during aging and become more densely packed on the dendrites of old bushy neurons. Alternatively, it is also possible that neurons with decreased dendritic synapses during aging were lost or in bad condition, and were not sampled in our data population. Taken together, the observed changes in bushy cell morphology and AN synapses may be an underestimation of neuronal modifications during aging in contribution to the development of ARHL.

## ACKNOWLEDGEMENTS

All images were generated at OSU Campus Microscopy and Imaging Facility (CMIF), supported by grant P30CA016058.

